# BOND-PEP: topology-conditioned bipartite alignment for evidence-grounded peptide binder generation

**DOI:** 10.64898/2026.02.18.706554

**Authors:** Wenze Ding

**Affiliations:** School of Mathematics and Statistics, Faculty of Science, University of Sydney, NSW 2006, NSW, Australia; Charles Perkins Centre, University of Sydney, Sydney, NSW 2006, Australia

**Keywords:** peptide design, sequence-first generation, retrieval-augmented generation, bipartite topology conditioning, evidence-grounded decoding

## Abstract

Peptide binders can modulate proteins that remain challenging for small molecules, but discovering high-affinity, selective peptides is still slow and sample-intensive. Sequence-first generators could scale design when structures are unavailable or conformationally heterogeneous, yet they often trade diversity for control: unconstrained sampling is inefficient while conditioning remains largely implicit. This limitation is exacerbated by the uneven transfer of protein language model priors to short peptides. Here we present BOND-PEP, a retrieval-augmented, bipartite-aligned, topology-conditioned framework that converts empirical binding evidence into an explicit, residue-resolved conditioning state for peptide generation. BOND-PEP shows low perplexity together with satisfactory free-generation hit rates and sequence novelty under a fair evaluation protocol and decoding budget. Compared with existing peptide generation methods, BOND-PEP achieves state-of-the-art results that match or improve upon validated peptide-protein sequence pairs. In total, BOND-PEP provides a practical, sequence-only route to controllable de novo peptide binder generation under noisy labels and distribution shift.

## Introduction

Peptide binders provide a direct route to modulate proteins that are difficult to drug with small molecules, including targets lacking well-defined pockets, exhibiting substantial conformational plasticity or containing intrinsically disordered regions^1-4^. Notably, estimates suggest that upwards of four in five disease-associated proteins remain undruggable with conventional small-molecule strategies, underscoring the scale of this unmet need^5-8^. Their therapeutic and functional potential spans competitive inhibition, induced proximity, and targeted degradation^9-11^. Yet the discovery of high-affinity, selective peptides remains labour-intensive and iterative^12-17^. From a learning and generalization perspective, peptide binder design is a conditional generation problem under severe constraints: only the target protein is consistently available; the design space grows exponentially with peptide length; training labels are sparse and noisy; and practical success requires robustness to distribution shift across protein families, binding modes, and disorder content^2, 3, 18-22^. Bridging this gap calls for generative methods that are not only expressive, but also controllable and stable under realistic data imperfections.

A dominant tradition in peptide binder design is structure-centric. Ranging from docking and Rosetta-style optimization to modern structure-conditioned neural generators, these approaches can directly enforce geometric complementarity and physicochemical constraints when reliable three-dimensional information is available^18, 19, 22-29^. However, structure-driven workflows inherit several practical bottlenecks: many targets lack high-confidence structures in relevant conformational states; binding often involves flexible or intrinsically disordered regions where a single static template is inadequate; errors in structure prediction or docking can be amplified during design; and the resulting multi-stage pipelines can be computationally heavy and brittle. These limitations motivate complementary sequence-first approaches that can operate when structure is missing, uncertain, or dynamically heterogeneous.

In parallel, large protein language models have enabled sequence-based representations with coevolutionary dependencies and residue-level constraints^30-35^. Embeddings with such signals have become a practical substrate that support peptide binder discovery and design without explicit structural templates. However, an often-overlooked gap is that protein-trained PLMs transfer unevenly across sequence regimes: they encode proteins well, yet can underperform markedly on short peptides.

Recent sequence-first methods fall broadly into two paradigms^2, 3, 36-39^. The first follows a “generate-then-rank” loop: a generator proposes many candidate peptides, and a contrastively learned compatibility model ranks and selects those most consistent with the target. This decoupling offers flexibility and can exploit large collections of unlabelled sequences, but the separation between generation and target–peptide alignment in the other hand leads to unstable search dynamics, where success depends on extensive sampling and strong post hoc filtering. The second paradigm performs direct target-conditioned generation by casting peptide design as conditional sequence infilling, where a masked binder span is reconstructed end-to-end from the target sequence. While this can improve coherence and reduce reliance on downstream filtering, the conditioning is frequently implicit, being encoded as general context rather than as explicit, fine-grained evidence of which residue-level patterns are preferred for a given target.

Consequently, existing sequence-first pipelines still struggle with the core tension between creativity and controllability, namely exploring novel sequences while remaining anchored to actionable binding evidence. What remains missing is a principled bridge from evidence to generation that simultaneously (i) grounds peptide priors in peptide-relevant statistics rather than assuming protein-derived priors transfer, and (ii) aligns target residues with candidate peptide patterns and exposes the resulting preferences as explicit conditioning.

Here we address this gap by introducing BOND-PEP, a retrieval-augmented, alignment-conditioned framework for peptide binder generation. BOND denotes Bipartite cONditioned Decoding, and BOND-PEP comprises three tightly coupled components. First, we retrieve a small set of candidate peptides from a large library to anchor generation in a locally relevant region of sequence space. This step injects historically plausible binding patterns as empirical prior knowledge to sharply reduce the effective search space to a target-relevant local manifold. Second, we perform residue-level bipartite alignment between the target protein and retrieved peptide candidates via a topology conditioner to distil token-level target-specific preference signals into an explicit conditioning state. More specifically, protein tokens attend to peptide tokens to identify which exemplar fragments are informative for the target, while peptide tokens attend back to protein residues to resolve where and how the target is likely to accommodate those patterns. Multi-round message passing integrates these cues into a final topologically aligned representation for later conditional decoding. Third, a conditional decoder generates novel peptides under this aligned evidence, effectively rewriting and recombining plausible motifs rather than sampling unconstrained sequences. This design encodes principled inductive biases: robust generation should begin from a locally relevant region of sequence space via retrieval and should be guided by residue-level correspondence signals from bipartite alignment, rather than by unconstrained sampling. Overall, by turning retrieved binding evidence into residue-resolved conditioning, BOND-PEP enables fully sequence-first, target-conditioned de novo peptide binder generation, providing a scalable route to therapeutic binders for protein targets that sample diverse conformations or contain substantial intrinsic disorder.

## Results and Discussion

### PLMs encode proteins well but underperform on peptides

Protein language models (PLMs) which learn transferable representations on large-scale protein sequences have become the most widely used foundation models for protein-sequence-related analysis and tasks. However, their performances on peptides have not been systematically evaluated. Here, we picked two widely adopted and state-of-the-art PLMs, ESM-2^31^ and more recent ESM-C^32^ and compared them across protein and peptide regimes. Stratifying sequences into proteins and peptides binned by length (>15, 10–15, ≤10 amino acids) with 1000 randomly sampled sequences per category, we evaluated these two models on three complementary sequence-modelling tasks: (i) self-copy (sequence self-consistency when the full sequence is provided), (ii) LOO (leave-one-out masked recovery, i.e., predicting one held-out residue at a time), and (iii) denoising (iterative denoising from partially corrupted inputs).

As shown in Fig.1., we found satisfactory performance on proteins but a pronounced degradation on peptides. Across tasks, proteins consistently formed a “high-performance regime”, whereas peptides—especially ≤10 amino acids —showed a sharp drop in recoverability and confidence. Meanwhile, these two PLMs showed complementary strengths. ESM-2 consistently outperformed ESM-C in the self-copy task. By contrast, ESM-C generally matched or exceeded ESM-2 when the input was partially observed.

**Fig. 1.**
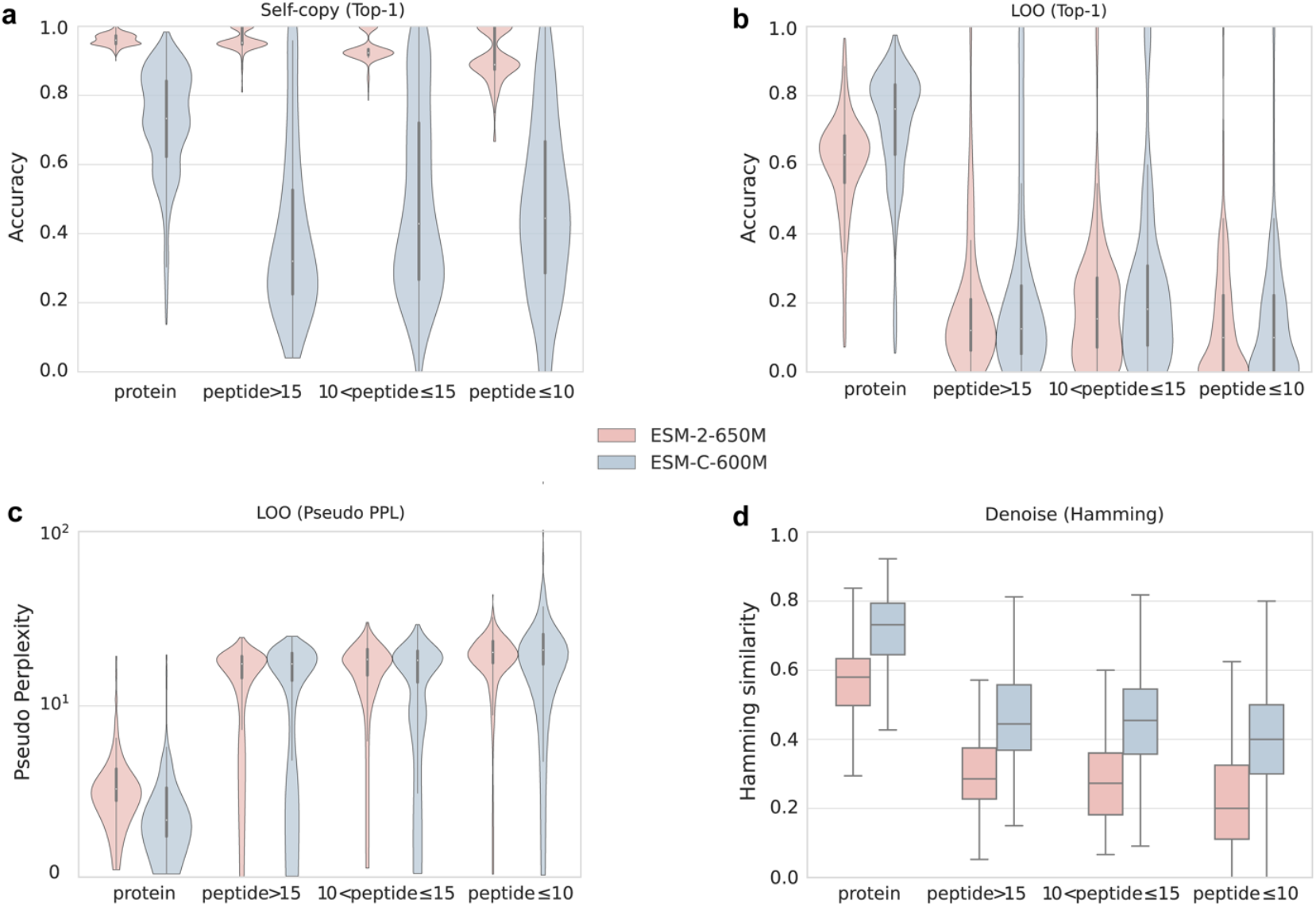
Protein language models exhibit a pronounced performance drop on peptides. We compared ESM-2-650M (pink) and ESM-C-600M (blue) on proteins and peptides stratified by length (protein; peptide: >15, 10–15 and ≤10 amino acids). For each category, 1000 sequences were randomly sampled from our sequence dataset. **a**. Self-copy (Top-1). Top-1 accuracy when the full sequence is provided as input and the model is evaluated on recovering the same residues. **b**. Leave-one-out (LOO; Top-1). Top-1 accuracy when one residue is masked at a time and predicted from the remaining context. **c**. LOO (pseudo perplexity). Predictive uncertainty under LOO, computed from per-position negative log-likelihood and exponentiated. Lower values indicate higher confidence. The y axis is log scale. **d**. Denoising (Hamming similarity). Reconstruction fidelity after iterative denoising from partially corrupted inputs, quantified as the fraction of positions matching the ground-truth sequence.

More specifically, in self-copy task, ESM-2 approached near-saturated Top-1 accuracy across all regimes, yet performance degraded on peptides, with the clearest decline for short peptides. In parallel, ESM-C remained strong on proteins but collapsed in performance on peptides. It showed uniformly lower accuracy, centred around 0.4, across peptide length bins (Fig. 1a). This peptide weakness became more notable under LOO task, which probes whether a model can infer a residue from its surrounding context rather than simply echoing it. Both models achieved markedly higher Top-1 LOO accuracy on proteins while performed poorly on peptides (Fig. 1b). Consistently, pseudo-perplexity of two models was lowest for proteins and substantially higher for peptides, with the shortest peptides showing the highest uncertainty (Fig. 1c). Finally, in denoising task, where sequences were corrupted and iteratively reconstructed, both models again showed a clear protein–peptide gap. Compared with proteins which were recovered with higher Hamming similarity, peptides exhibited reduced reconstruction fidelity, and the deficit increased with decreasing peptide length (Fig. 1d).

Together, these results show that the statistical regularities learned from proteins transfer poorly to short peptides, no matter the ground-truth context is fully visible or partially masked. This hints that the limited context of short peptides provides much fewer informative constraints for each position, and thus further reduces residue identifiability and force the models to distribute probability mass across multiple plausible amino acids. Because of the intrinsic difference in sequence length, short peptides sit outside the “comfortable zone” that these PLMs implicitly rely on. This gap motivates downstream analyses of peptide representation space and the development of peptide retrieval and generation strategies.

### Contrastive retrieval de-collapses peptide representations

We found that ESM-2 peptide embeddings undergo a marked collapse, rendering distinct peptides nearly indistinguishable in representation space. In the global ESM embedding space, peptide representations concentrate tightly along the dominant principal component, whereas protein representations remain broadly distributed (Fig. 3a). This compressed geometry implies that local operations in ESM space of peptides, such as nearest-neighbour search or embedding-space perturbation, will be poorly aligned with meaningful biological variation and instead behave as quasi-random jitter within that collapsed region of the space.

We therefore examined nearest-neighbour search as the most direct test of locality in the ESM peptide embedding space. We randomly selected 200 protein-peptide pairs in our dataset, and then embedded the target proteins and all library peptides with ESM. For each protein, we denote its paired peptide as the ground-truth (GT) peptide. We then retrieved the top-256 peptides from the library as its candidate set (Top) by ranking peptides by cosine similarity to the protein embedding. As a control, we also sampled an equally sized peptide set uniformly at random (Random).

We found that candidate-set intra-similarity was extremely high for both Top and Random sets (0.9102 ± 0.0365 vs 0.9349 ± 0.0058, Fig. 3b, e), consistent with peptides occupying a narrowly concentrated region of the ESM embedding space irrespective of selection strategy. More strikingly, the Top retrieval did not enrich proximity to the GT peptide. The mean cosine similarity to GT was even lower for the Top set than the Random (0.8615 ± 0.0422 vs 0.9421 ± 0.0213). Likewise, the overlaps between the candidate sets and the GT KNN neighbourhood (the 256 nearest peptide neighbours of the GT peptide in ESM space) were both near zero (Fig. 3e, Fig. S1-S2). The lack of enrichment was also reflected at the sequence level for both Jaccard similarity and BLOSUM62-based similarity (Table S1-S3). Together, these results indicate that distances in the raw peptide embedding space are not operationally meaningful under such collapse.

Despite the collapse of raw peptide geometry in ESM space, binding-pair supervision can still yield effective discriminators for protein–peptide matching, as demonstrated by CLIP-style dual-encoder models^2^. Yet, it remains unclear how such supervision reshapes peptide representations and whether the resulting geometry supports scalable library indexing and fast querying. We therefore developed a target-conditioned peptide retriever (Fig. 2a) to interrogate what binding supervision is learning.

**Fig. 2.**
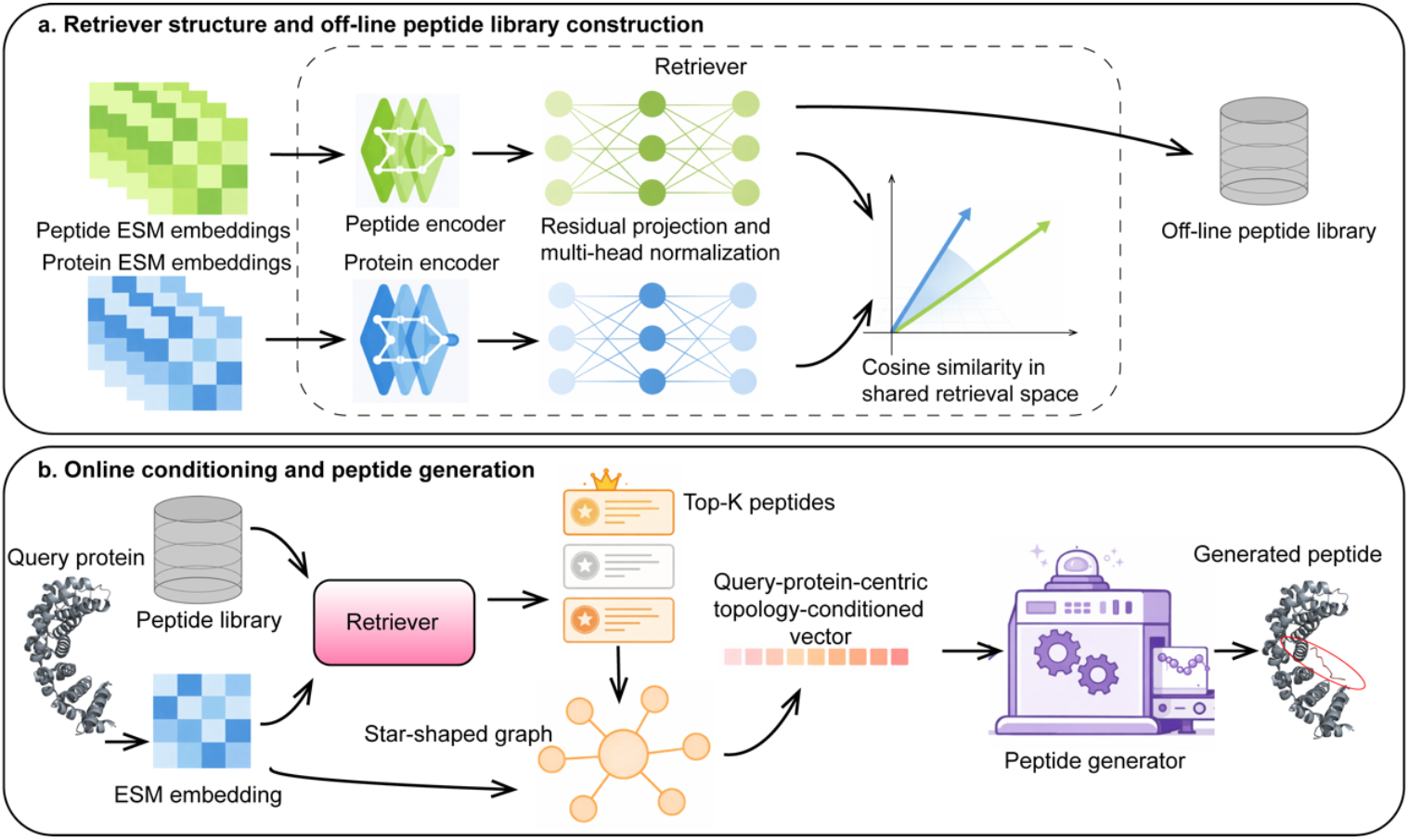
Target-conditioned peptide retrieval and topology-conditioned peptide generation. **a**. Retriever architecture and off-line peptide library construction. ESM-derived embeddings of peptides and proteins are fed into peptide and protein encoders, respectively, followed by a residual projection module with multi-head normalization to map both modalities into a shared retrieval space. Candidate peptides are then ranked by cosine similarity between the query protein and peptide representations. Peptide embeddings for the entire library are precomputed and indexed off-line to enable fast search at query time. **b**. Online conditioning and peptide generation. Given a query protein, its embedding is used to retrieve the top peptides from the pre-prepared indexed library. These retrieved peptides are used to construct a protein-centred star graph for message passing. A query-centric, topology-conditioned vector is then derived and provided as conditioning input to a peptide generator to produce new peptide. The binding pose of the generated peptide to the query protein is highlighted at right with a red ellipse.

**Fig. 3.**
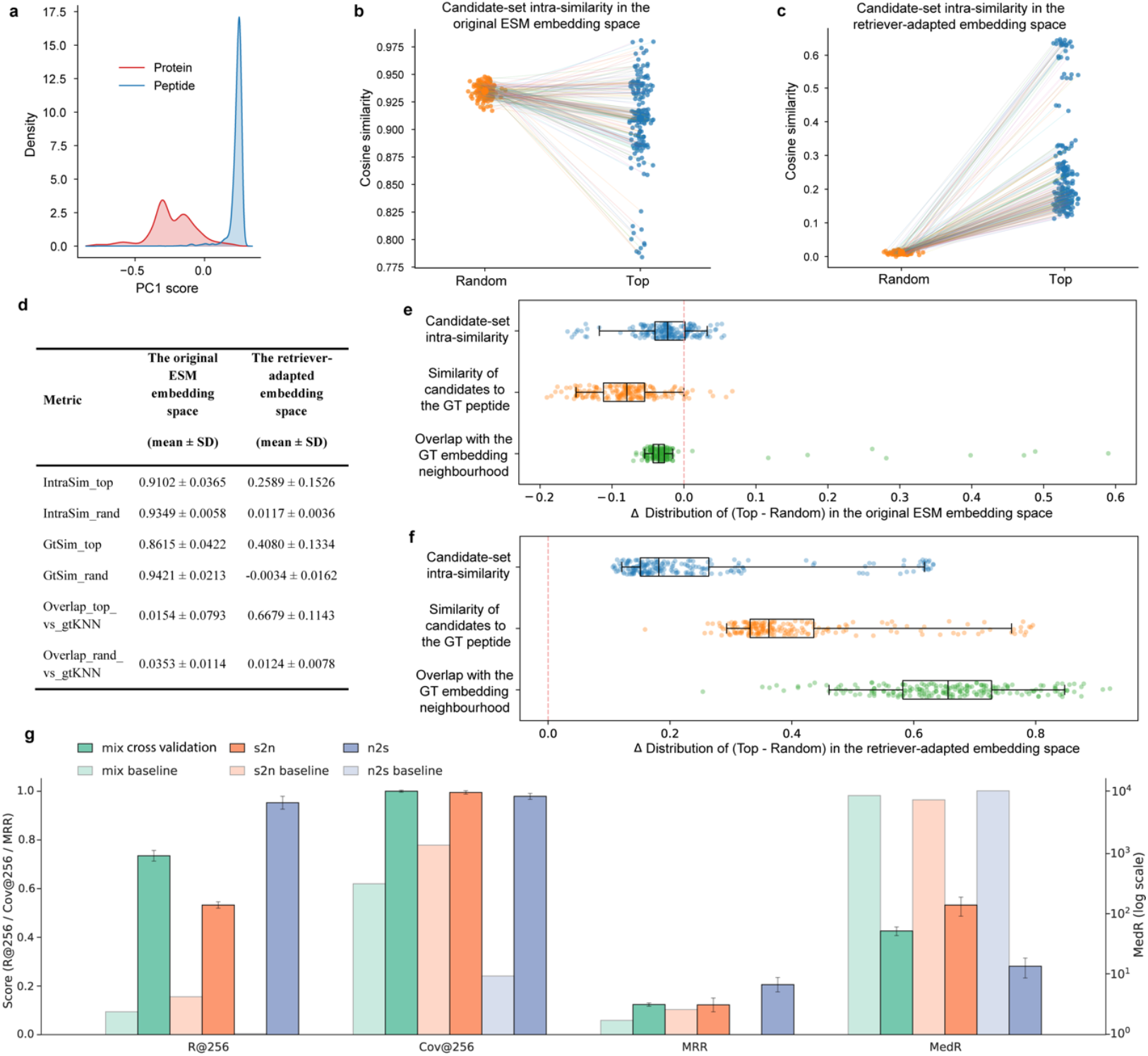
Original ESM peptide embeddings collapse in geometry, whereas binding supervision restores target-aligned local structure for retrieval. **a**. Distribution of first principal component (PC1) scores of ESM embeddings for proteins and peptides. Peptide embeddings concentrate along the dominant component, whereas protein embeddings remain broadly distributed. **b-c**. Candidate-set intra-similarity (mean pairwise cosine similarity within the set) for K = 256 peptides per query in the original ESM embedding space (b) and in the retriever-adapted embedding space (c). For each query protein (n = 200), Top denotes the top-K peptides ranked by cosine similarity between the query-protein embedding and peptide embeddings from the whole peptide library, and Random denotes a size-matched uniformly sampled peptide set. Points represent individual queries; lines connect paired Random and Top sets. **d**. Summary of locality metrics in the original ESM and retriever-adapted spaces. Metrics are computed per query protein using K = 256 peptides. IntraSim_top and IntraSim_rand denote the mean pairwise cosine similarity among peptides within the Top and Random candidate sets, respectively. GtSim_top and GtSim_rand denote the mean cosine similarity between each candidate peptide in the Top or Random set and the paired ground-truth (GT) peptide embedding. Overlap_top_vs_gtKNN and Overlap_rand_vs_gtKNN denote the overlap between the candidate set (Top or Random) and the GT neighbourhood (the K nearest peptide neighbours of the GT peptide in the corresponding embedding space). **e-f**. Per-query differences (Top − Random) for the three locality metrics in the original ESM space (E) and the retriever-adapted space (F). Each point represents one query protein. Boxplots show the median and interquartile range. Red dashed lines indicate zero difference. **g**. Performance of our target-conditioned retriever and an ESM baseline across three evaluation protocols: n2s (train on noisy, test on strict), s2n (train on strict, test on noisy) and mix (noisy + strict with cross-validation). Recall@256, Coverage@256 and mean reciprocal rank (MRR) are plotted on the left y axis (range 0-1 and higher is better), whereas median rank (MedR) is plotted on the right axis (log scale and lower is better). Bars show the mean across 10 repeats and error bars indicate s.d.

Across evaluation protocols, our retriever showed robust and practically useful performance, consistently outperforming the ESM baseline (Fig. 3g). We evaluated three settings: n2s (trained on the noisy dataset and tested on strict); s2n (the reverse direction, i.e., train strict, test noisy) and mix (merging the noisy and strict datasets followed by cross-validation) and each setting was repeated 10 times. Performance was assessed using standard information-retrieval metrics: Recall (GT recovered within the top retrieved candidates), Coverage (library-level breath, defined as the fraction of unique peptides appearing in the union of all top retrieved candidate lists across queries), MRR (mean reciprocal rank), and MedR (median rank, lower is better). Because the noisy and strict datasets are constructed using different filtering criteria, these protocols impose different degrees of distribution shift and difficulty: n2s is comparatively easier, whereas s2n is the most challenging. Nevertheless, our retriever maintained strong performance across all settings, supporting that it captures generalized structure across distributions. We further examined the sensitivity of performance to the choice of retrieval depth K (Fig. S3). Balancing retrieval quality against downstream computational cost, we selected K as 256.

We then asked what binding supervision changes in representation space. To test whether performance reflects improved local geometry rather than scoring alone, we repeated our locality analysis described above in the retriever-adapted embedding space. The results revealed a pronounced geometric “de-collapse”. In the retriever-adapted embedding space, Random sets became effectively unstructured, while Top sets formed coherent, query-specific neighbourhoods, with intra-set similarity as 0.0117 ± 0.0036 vs 0.2589 ± 0.1526 (Fig. 3c, f). Crucially, this coherence aligned with ground truth. Top candidates showed substantially higher similarity to the GT peptide than Random (0.4080 ± 0.1334 vs −0.0034 ± 0.0162, paired Wilcoxon P = 1.44×10^−34^, Fig. 3f, Fig. S1) and recovered the GT local neighbourhood with significantly higher overlap (0.6679 ± 0.1143 vs 0.0124 ± 0.0078, P = 1.43×10^−34^, Fig. 3f, Fig. S2). This improvement was also reflected at the sequence level, arguing against a purely geometric artefact. Retrieved Top peptides exhibited significantly higher internal sequence coherence than Random and substantially greater sequence similarity to the GT peptide (Table S1-S3). Together, these results show that our retriever expands the effective volume of peptide representations and restores discriminative, target-aligned local structure, enabling scalable offline indexing and fast online querying.

### Topology-conditioned bipartite alignment enables evidence-grounded and controllable decoding

We first examined the organization of peptide binders in the retriever embedding space. For a given target protein, experimentally supported binders concentrate in a local neighbourhood around the target representation (Fig. 4a), suggesting that binding-competent sequences occupy a target-specific local manifold with similar sequence patterns and physicochemical profiles rather than being uniformly distributed in sequence space. Consistent with this, multiple distinct peptides can bind the same target protein, as illustrated by Fig. 4b. Together, these observations motivate a generation strategy that anchors decoding to locally relevant evidence, while still allowing controlled departures from any single retrieved exemplar.

**Fig. 4.**
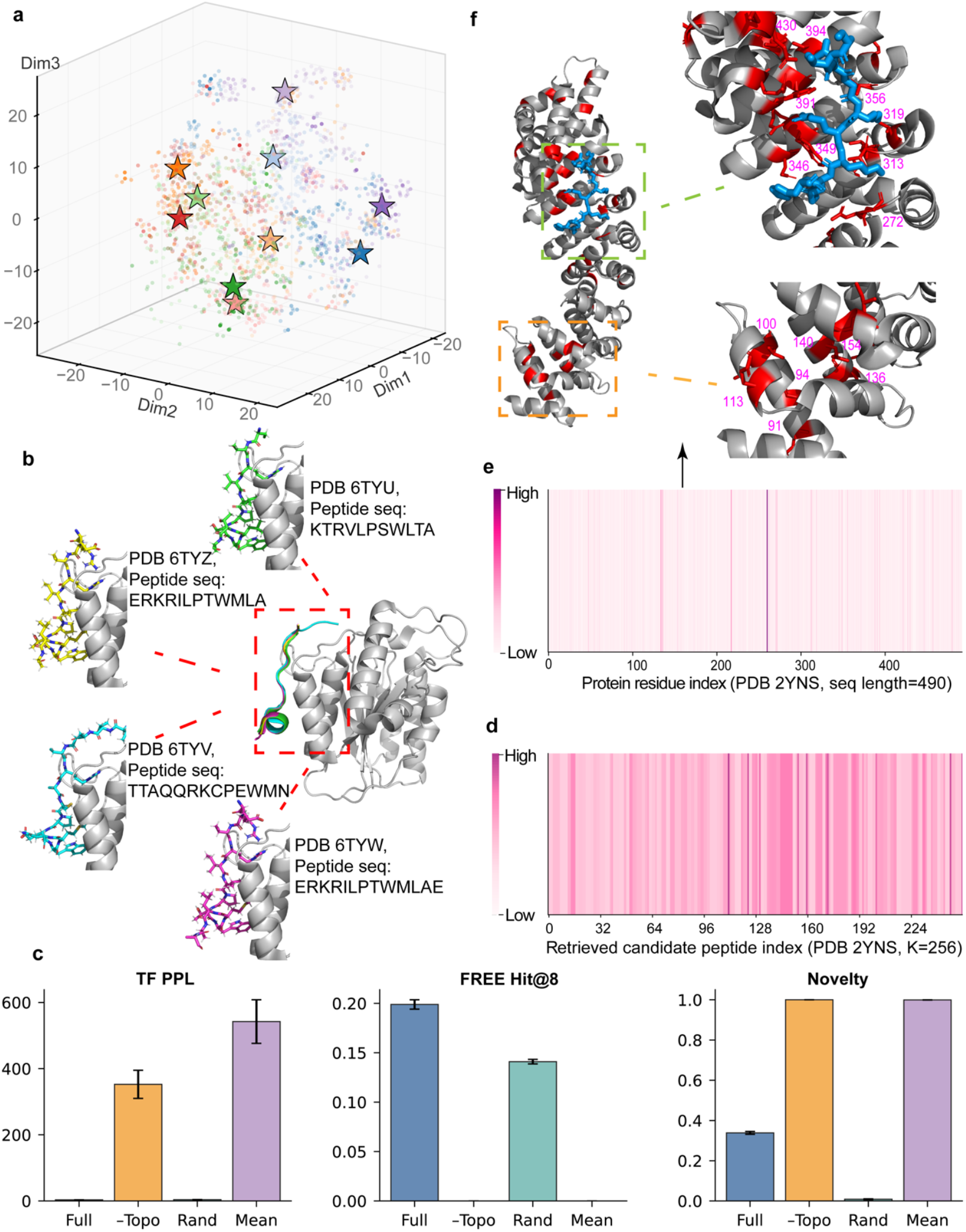
Topology-conditioned bipartite alignment yields residue-level preference states for evidence-grounded peptide decoding. **a**, 3D t-SNE visualization of the frozen retriever embedding space for proteins (stars) and peptides (dots). For each protein target, associated peptides (same colour) occupy a local neighbourhood around the target representation. **b**, A single target can bind multiple distinct peptides, illustrated by four complexes with the same receptor fold and different bound peptides (PDB 6TYU/6TYV/6TYW/6TYZ). Receptor is coloured by grey cartoon. Zoomed views highlight the binding site, where different peptides are shown as sticks in different colours with sequences annotated. **c**, Ablation of topology conditioning. Teacher-forcing perplexity (TF PPL), free-generation hit rate among the top 8 generated peptides (FREE Hit@8) and novelty for the full model (“Full”) versus removing the topology conditioner (“–Topo”), using a random selected context instead of the retrieved one (“Rand”) or using mean-pooling in place of the topology conditioner (“Mean”). All comparisons use identical training and decoding settings, and metrics are reported at the final training epoch (epoch 70) for each variant. Error bars indicate variability across independent runs. **d**, Preference over retrieved peptides (K=256) for an example target (PDB 2YNS), indicating distributed evidence usage rather than dependence on few exemplars. **e**, Protein-side residue preference along the 2YNS sequence (length 490), revealing sparse high-preference positions. **f**, Structural mapping of high-preference residues (red) onto the receptor (grey) with the bound peptide (blue). The interface-proximal region (dashed green outline) and a distal region (dashed yellow outline) are highlighted and shown as zoomed insets on the right, where magenta labels denote residue indices.

We therefore instantiate this concept as a topology-conditioned bipartite alignment module that converts the retrieved peptides into an explicit, protein-centric conditioning state (Fig. 2b). The key design choice is to not treat these peptide candidates as independent “prompts”, but to couple them with the query protein through a local protein-peptide bipartite context. This topology enforces a protein-centric view: peptides do not communicate with one another and subgraphs are isolated per protein, so evidence integration is mediated exclusively by the query protein. We then perform bidirectional message passing over this bipartite structure to distil two complementary signals that retrieval alone cannot provide: (i) which candidates are informative for the current target (from peptide-to-protein aggregation), and (ii) where on the protein these candidates consistently constrain compatibility (from protein side). Finally, we inject this topology-conditioned state into a Transformer decoder as memory token, enabling evidence-grounded decoding under a stable, target-specific conditioning signal.

We next asked whether topology conditioning is necessary for practical peptide sequence generation. We performed a controlled ablation that isolates the effect of the topology-conditioned alignment while keeping the same retriever, decoding budget, and evaluation protocol. As shown in Fig. 4c, removing topology conditioning (“–Topo”) led to a pronounced degradation in generation quality compared with the full model (“Full”): teacher-forcing perplexity increased sharply and FREE Hit@8 collapsed to near zero, indicating that retrieval alone is insufficient unless it is converted into a usable conditioning state for the generator. Then, we replaced the retrieved context with a random set of peptides but kept the architecture of topology conditioner (“Rand”). Compared with the full model, this setting partially preserved Hit@8, but its novelty dropped markedly, consistent with a failure mode where the model relies on superficial sequence reuse rather than target-specific preference shaping. Finally, we kept the retrieved context but replaced the topology conditioner with mean pooling aggregation (“Mean”). We found this setting also failed to provide a functional conditioning signal, again showing high perplexity and negligible Hit@8. Additional supporting data and complementary analyses are provided in Fig. S4 and Tables S4–S5. Together, these results indicate that topology conditioning provides a strong inductive bias, steering generation toward target-compatible sequence motifs. This focuses probability mass on productive regions of sequence space, improving sample efficiency and mitigating trivial copying of retrieved peptides.

To interpret the representations learned by the topology-conditioned alignment module, we visualized attention-based preference maps spanning retrieved candidates and protein residues. First, as shown by Fig. 4d, the model does not rely on a single retrieved peptide as evidence. Instead, attention over the K=256 candidate peptides exhibits a distributed usage pattern, supporting an evidence-grounded mechanism that integrates signals across multiple candidates rather than memorizing one exemplar. Second, the induced preference over the protein sequence is sparse and structured. When we summarize residue-level conditioning signals along the protein sequence, the preference profile shows distinct peaks at a subset of residues (Fig. 4e), indicating that the alignment focuses on specific positions that are repeatedly informative under different retrieved-context views. Mapping these high-preference residues back to the 3D structure reveals two complementary categories of sites (Fig. 4f). One group localizes to the peptide-binding region and lines the interaction surface (e.g., hotspots marked at the right top of Fig. 4f). The other group localizes to interior regions that are not directly at the interface but plausibly constrain the receptor geometry and stabilize the binding-competent conformation (hotspots marked at the right bottom of Fig. 4f).

Collectively, these visualizations clarify how topology-conditioned bipartite alignment operates. The module aggregates evidence from multiple retrieved peptides into a protein-centric, residue-resolved preference map, highlighting residues that likely mediate binding as well as positions that stabilize the bound conformation. This interpretable preference state can then be used to steer decoding, anchoring generation to empirically supported motifs while still permitting sequences that generalize beyond the retrieved candidates.

## Methods

### PLMs Evaluation

Evolutionary Scale Modeling (ESM) is a series of transformer protein language models developed by Meta AI’s FAIR team. Among them, ESM-2 (released in 2022) has become one of the most widely used PLMs in protein informatics. At the end of 2024, EvolutionaryScale introduced ESM-3 and ESM Cambrian (ESM-C) as parallel model families. While ESM-3 is designed as a protein generative model, ESM-C is oriented toward sequence representation learning aimed at underlying biology. Here, we evaluated ESM-2-650M and ESM-C-600M under identical preprocessing. Both models used the same standard amino-acid tokenizer, and we used the model’s native output logits directly, ensuring our metrics reflect each model’s intrinsic residue-level predictive distribution properly.

We stratified sequences into proteins and peptides grouped by length (>15, 10–15, and ≤10 amino acids), and then randomly sampled 1000 sequences for each category from our sequence dataset. To evaluated their performance gap between proteins and peptides, ww run ESM-2-650M and ESM-C-600M on three complementary tasks: self-copy, LOO and denoising.

For self-copy, the full unmasked sequence *s* = (*y*_1_, …, *y*_*L*_) was provided as input, and the model produced logits at each position. We computed per-position Top-k correctness by checking whether the true amino acid *y*_*i*_ appeared among the k highest-probability tokens at position *i*. We reported Top-1 accuracy in our results. Sequence-level accuracy was the mean across positions as following:

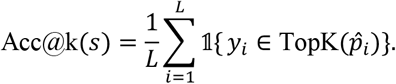

For LOO, we evaluated each position independently by masking position *i* and predicting *y*_*i*_ from the remaining context. LOO task is more stringent than self-copy, and we reported not only Top-1 accuracy but also pseudo-perplexity (pPPL) to quantify overall predictive uncertainty. pPPL is derived from the negative log-likelihood (NLL). Averaging NLL over positions gives the typical “surprisal” of native residues from a sequence. Exponentiating this quantity yields pPPL, which can be interpreted as the model’s effective number of competing choices per position. Lower pPPL means less uncertainty, which further means the better model performance. We computed pPPL via:

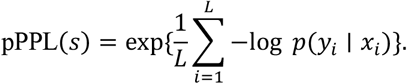

For denoising, we first corrupted the sequence by randomly replacing 20% residues and then iteratively refined the sequence for *n* steps. *n* equals the number of replaced residues. For each refinement step, the lowest-confidence position under the current model prediction was identified and then resampled via masked inference. We used Hamming similarity to quantify the reconstruction quality between the final sequence *ŝ* and the ground truth *s* as following:

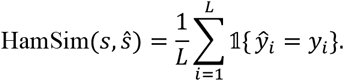

### Network Architecture Overview

We developed a retriever-conditioned, and topology-aware autoregressive peptide generator. Given a query protein, the model would (i) encode the protein into a dense vector, (ii) retrieve a context set of *K* candidate peptides from a large peptide library using a frozen dual-encoder retriever, (iii) construct a local bipartite subgraph between the query protein and the retrieved peptides, (iv) perform multi-layer bipartite message passing to produce a topology-conditioned protein embedding, and (v) left-to-right generate candidate peptide sequence. Training examples consisted of protein–peptide pairs, and we used five-fold cross-validation with shuffling throughout. Unless otherwise stated, all architectural choices and hyperparameters were selected based on cross-validation.

### Precomputed token embeddings

All proteins and peptides were represented using precomputed token-level embeddings from ESM-2-650M, stored on CPU to minimize GPU memory usage. Specifically, we stored: protein embeddings 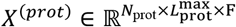 and corresponding masks 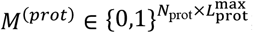, peptide embeddings 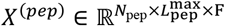 and corresponding masks 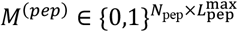. Here 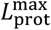 and 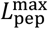 denote the maximum sequence length within the protein and peptide libraries. We padded proteins and peptides to these fixed max lengths respectively and indicated padded positions by mask value 0. *F* is the per-token embedding dimension produced by ESM-2-650M. It is worth noting that during network computation, only the embeddings required for a mini batch were transferred to GPU using non-blocking copies.

We treated the ESM special BOS token differently for proteins and peptides. For peptides, we discarded BOS, thus each peptide mask contains exactly *L* ones corresponding to residues. In contrast, for proteins we retained BOS, resulting in 1 + *L* valid positions per protein sequence. This choice was determined by strictly controlled comparison in which we evaluated models trained with and without BOS token for each modality and selected the configuration that yielded the best performance. We hypothesize that BOS is less informative for short peptides, consistent with the degradation of peptide representations observed in our previous analyses, whereas for full-length proteins the BOS token captures useful global context.

### Sequence encoders

We use an attention module called AttnSeqEncoder as our protein encoder. Given a batch of ESM protein token embeddings 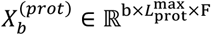 and corresponding masks, we first project each token to a hidden representation: *h*_*t*_ = *f*_tok_(*x*_*t*_) ∈ ℝ^*d*^, where *f*_tok_ is a multi-layer perceptron (MLP) with layer normalization and dropout. From this, we compute scalar attention logits using: *e*_*t*_ = *w*^⊤^ tanh(*Wh*_*t*_) ∈ ℝ. We mask invalid tokens by setting logits to −∞, and thus form normalized attention weights:

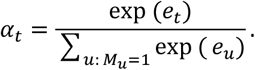

Finally, the pooled protein embedding is obtained by applying a linear head to the attention-weighted sum of token representations, yielding a *D*-dimensional vector (default *D* = 512):

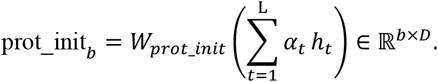

Analogously, peptide sequences are encoded with a convolutional module called ConvSeqEncoder. First, a batch of ESM token embeddings of peptides 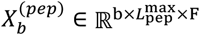 with corresponding masks are transposed and projected into a model width *d*_model_. Then a stack of residual 1D convolution blocks are applied as:

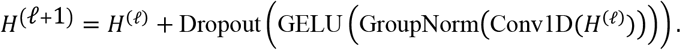

The final pooled peptide embedding is computed by applying a linear head to the masked mean pooling:

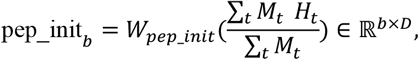

### Retriever architecture

The retriever is implemented as an independent dual-encoder module that embeds proteins and peptides separately into a shared retrieval space, enabling efficient maximum-inner-product search for any query protein over large peptide libraries. Given token-level embeddings and masks, the protein tower (AttnSeqEncoder) and the peptide tower (ConvSeqEncoder) produce fixed-dimensional representations prot_init, pep_init ∈ ℝ^*D*^, which are then mapped into the shared retrieval space by modality-specific projection heads:

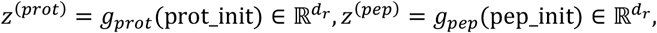

where *d*_*r*_ = 2048 is the dimension of the retrieval space. Both projection heads *g*_*prot*_(⋅ ) and *g*_*pep*_(⋅) are implemented as residual MLPs with LayerNorm and Dropout:

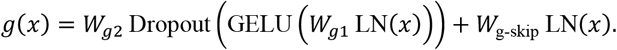

This design improves optimization stability compared to a single linear projection, while preserving a direct path from normalized base embeddings to the retrieval space.

Candidate peptides are then ranked by dot-product similarity:

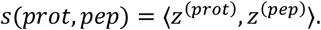

This is equivalent to cosine similarity when using embedding normalization. The factorized scoring allows offline precomputation of peptide embeddings and scalable top-*K* retrieval without cross-attention between modalities. To stabilize similarity score *s*(*prot, pep*) in the high-dimensional retrieval space, we normalize vectors using a multi-subspace scheme. More specifically, for any projected vector 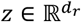, we reshape it into *M* sub-vectors 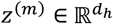 where *d*_*h*_ = *d*_*r*_/*M*, then normalize each sub-vector independently:

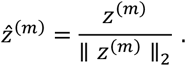

We further introduce learnable head importances via logits *γ* ∈ ℝ^*M*^, forming weights

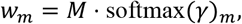

and apply them as 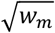 scaling:

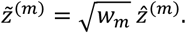

Finally, we scale by 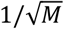 and concatenate:

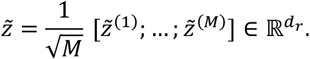

With this construction, the dot product decomposes into a weighted sum of per-subspace cosines:

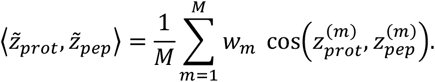

This multi-subspace or multi-head scheme yields stable magnitudes and allows the model to learn which subspaces are more informative for retrieval.

### Retriever pretraining

We pretrain the retriever using a two-stage contrastive objective. Specifically, we optimize the retriever encoders, projection heads, and head-importance logits *γ* by minimizing a weighted sum of an in-batch contrastive loss and an auxiliary hard-negative mining loss:

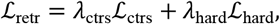

with *λ*_ctrs_ = 1.0 and *λ*_hard_ = 0.3. To make the training stable, hard-negative mining is enabled only after the 12th epoch.

Given a mini-batch of *b* (default batch size *b* is 512) matched protein–peptide pairs 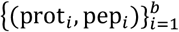, we compute normalized retrieval vectors 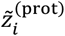 and 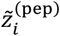 as described above, and then form a temperature-scaled similarity matrix:

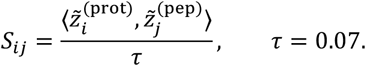

Using the paired examples (*i, i*) as positives and all off-diagonal entries as in-batch negatives, we minimize a cross-entropy loss for protein→peptide retrieval:

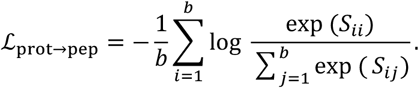

To avoid modality-specific collapse and enforce bidirectional alignment, we use the symmetric variant by adding the reverse direction, *i*.*e*., peptide→protein retrieval:

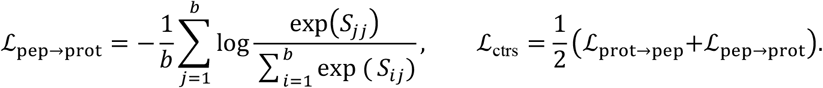

To further sharpen the retrieval beyond in-batch negatives, we additionally optimize a hard-negative objective after the 12th epoch. At initialization and after every epoch, we refresh a CPU-resident peptide index by encoding the full peptide library with the current retriever parameters to obtain a normalized retrieval matrix

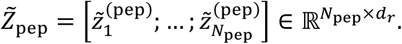

For a subset of query proteins per training step, we retrieve 2048 peptides by dot-product search against 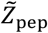. To reduce ambiguity from near-duplicates or extremely similar candidates, we discard the highest-ranked 64 retrieved peptides and select the subsequent 256 candidates as hard negatives 𝒩_*i*_. We then increase separation between the positive peptide and the mined negatives using a contrastive loss of the following form:

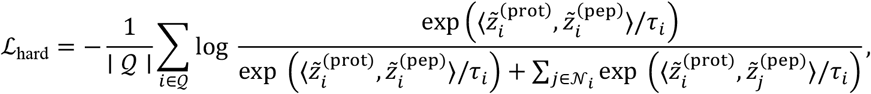

where *τ*_*i*_ = max (*τ, τ*_min_) applies a temperature floor *τ*_min_ = hard_temp_floor = 0.1 for numerical stability. This auxiliary term explicitly trains the retriever to down-rank highly confounding peptides that are close to the query in the current embedding space.

We optimize retriever parameters using AdamW with learning rate lr = 4 × 10^−4^ and weight decay weight_decay = 10^−4^. A stepwise learning-rate schedule is applied. To select retriever checkpoints robustly, we performe 5-fold cross-validation with shuffling. For evaluation, we compute Recall@ *K* for multiple *K* values (128, 256, 512 and 1024) on the held-out fold:

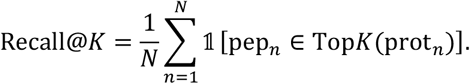

We also monitor the retriever via coverage of the peptide library defined as the fraction of unique peptides appearing in retrieved lists:

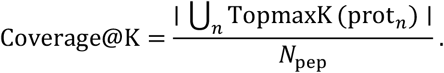

### ESM-embedding-space distribution analysis

To assess distributional differences between ESM representations of proteins and peptides, we randomly drew 2,000 protein–peptide pairs from our dataset. For each protein and peptide, variable-length ESM embeddings were converted to fixed-dimensional vectors by mean pooling. We first projected pooled embeddings to one dimension using principal component analysis (PCA) and then visualized distributional differences via kernel density estimation (KDE). Specifically, we estimated the marginal density of the first principal component for proteins and peptides separately using Gaussian KDE and plotted the resulting density curves as filled overlays. Density curves are normalized such that the area under each curve integrates to 1.

### Candidate-set level assessment of peptide-space

We performed a candidate-set level analysis on *N*_*q*_ = 200 randomly sampled proteins to probe how peptide representations differ between the original ESM embedding space and the retriever-adapted embedding space. For each protein query, we constructed a top-ranked candidate set under each space and compared it to a size-matched random baseline using paired measurements (top vs random for the same query). This design allowed us to quantify how each embedding space shapes (i) within-set cohesion, (ii) proximity to the ground-truth (GT) peptide, and (iii) overlap with the GT peptide’s local neighbourhood. We also did complementary sequence-based analyses to test whether candidates selected in the native ESM space versus the retriever-adapted space differ at the sequence level, including k-mer Jaccard and BLOSUM62 alignment similarity as a proxy for amino-acid substitution and biochemical property similarity.

More specifically, each entry is a tuple (*p*_*gt*_, *t*) specifying a protein indexed *t* and its associated GT peptide indexed *p*_*gt*_with amino-acid sequence string 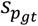. For each query protein *t*, we constructed two candidate peptide sets of size K = 256 from peptide library *p* ∈ 𝒫 in parallel: top candidate set 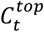 and random baseline 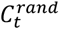 for both embedding spaces. For ESM embedding space, we first obtained protein and peptide representations 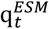 and 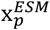 by mean pooling of ESM embeddings and then performed retrieval as:

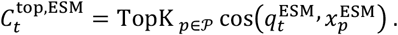

Meanwhile, for retriever-adapted embedding space, we loaded a pre-trained retriever checkpoint to compute representations of query protein 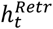 and peptide library 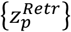. The retrieval was then performed by the retriever’s scoring function, yielding 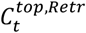. Their random baseline 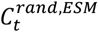 and 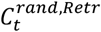 were sampled uniformly without replacement from the peptide library in corresponding embedding spaces.

All embedding-space metrics used cosine similarity. For each query (*p*_*gt*_, *t*), we computed the following metrics for both 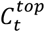 and 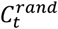 across all embedding spaces:

1. Candidate-set intra-similarity (within-set cohesion):

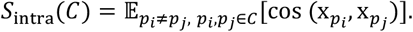

We approximated this expectation by sampling 4000 random ordered pairs (*i, j*) with *i* ≠ *j* per query.
2. Similarity of candidates to the GT peptide:

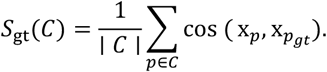
3. Overlap with the GT embedding neighbourhood: We first computed the GT peptide’s K-nearest neighbours in the same embedding space across the full peptide library:

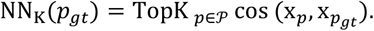

We then computed the overlap fraction:

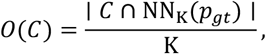

which captures whether the candidate set is enriched for peptides in the immediate embedding neighbourhood of the GT peptide. We also did complementary sequence-based analyses. Related sequence-space metrics are:
4. k-mer Jaccard similarity (*k* = 3): Given a peptide sequence *s*, we represent it by the set of all overlapping *k*-mers, denoted 𝒦_*k*_(*s*). For two sequences *a* and *b*, their *k*-mer Jaccard similarity is defined as:

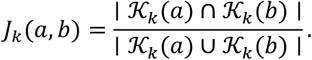

We then computed intra-set similarity and GT similarity of peptide sequences like (1) and (2).
5. BLOSUM62 alignment-based similarity: We computed global alignment scores under BLOSUM62 substitution scores using a Needleman–Wunsch dynamic program with affine gap penalties:

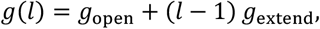

where *g*(*ℓ*) is the total penalty for a contiguous gap of length *ℓ, g*_open_ penalizes opening a new gap, and *g*_extend_ penalizes extending an existing gap by one position. We set *g*_open_ = −11 and *g*_extend_ = −1. For sequences *a, b*, let *S*(*a, b*) be the optimal global alignment score. To normalize for length and composition, we used:

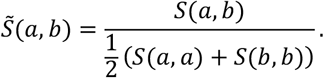

Similarly, intra-set similarity and GT similarity was computed like (4) as well.

For each metric *m*, we obtained paired per-entry measurements 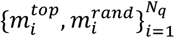 and analysed paired differences 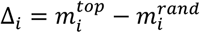. We assessed significance using two-sided Wilcoxon signed-rank test and paired t-test, quantifying effect size using paired Cohen’s *d* = mean(Δ)/sd(Δ).

### Retriever freezing and peptide library preparation

Once the training of retriever is completed, we load the pretrained retriever checkpoint and freeze all retriever parameters 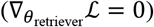, ensuring retrieval behavior remains fixed during all subsequent training and inference stages. To accelerate training and inference, we precompute retrieval embeddings for the entire peptide library. First, the peptide tower produces base embeddings pep_init_*i*_ ∈ ℝ^*D*^ for all peptides *i* ∈ {1, …, *N*_pep_}. Then the projection head and multi-subspace normalization produce retrieval embeddings 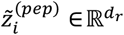. The resulting library matrix as following:

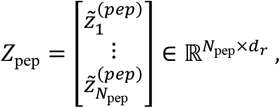

is stored on CPU to minimize GPU memory usage.

For each query protein, we compute the protein retrieval embedding 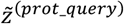 from the protein tower output and projection head. We then compute similarities by multiplication:

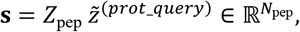

and select the indices of largest 256 scores. These indices define the candidate peptide set used downstream for topology conditioning and sequence generation.

### Topology conditioner

Topology conditioning is performed via a local bipartite subgraph per protein. For each mini-batch of size *b*, we construct this protein–peptide local bipartite graph *G* = (*V, E*) that couples each query protein to its retrieved top-*K* (default *K* = 256) peptide context. The node set is defined as:

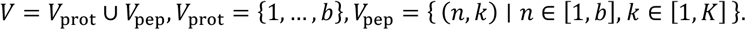

We instantiate two directed edge sets that form a star-shaped subgraph for each protein:

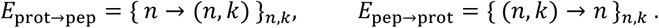

Thus, the graph contains exactly *bK* directed edges in each direction and no cross-protein edges among peptide candidates. Initial node features are given by the batch protein embedding prot_init_*n*_ ∈ ℝ^*D*^ and the retrieved peptide embeddings pep_init_sub_*n,k*_ ∈ ℝ^*D*^.

Topology conditioning is performed using *L* (default *L* = 3) stacked multi-head bipartite graph-attention layers (called MultiHeadBiGAT module). Each layer contains two directional multi-head attention blocks with independent parameters: (i) a protein→peptide update that aggregates protein information into each peptide node, and (ii) a peptide→protein update that aggregates peptide information into each protein node. For better description here, we use *H* denote the number of attention heads (default *H* = 4). For a generic directed update from source node *i* to destination node *j*, we apply pre-normalization (LayerNorm) and head-specific linear projections:

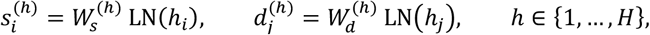

where *s* represents source and *d* for destination. Attention logits follow the additive form implemented in MultiHeadGAT and we also add a cosine-similarity bonus. Thus, the unnormalized edge score would be:

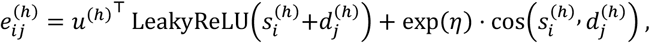

where *u*^(*h*)^ is a learnable head-specific vector and *η* is a learnable scalar. Normalized attention coefficients are computed for each destination node using a stable segment-wise softmax over its incoming edges followed by attention dropout:

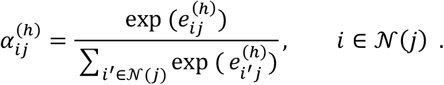

Messages are aggregated per head as a weighted sum of source projections:

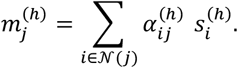

Head outputs are merged either by mean or concatenation, and then projected to the output width and combined with a residual connection from the destination state as following:

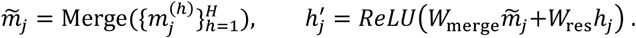

Applying this operator to both directions yields one bipartite layer update:

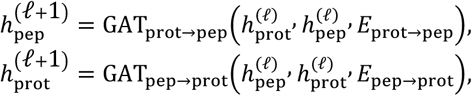

and we have *L* stacked layers as mentioned before.

After *L* bipartite layers, we take the updated protein node state as the final topology-conditioned vector. Notably, the vector remains protein-centric, while being selectively modulated by retrieved peptides through the peptide→protein attention. This attention mechanism learns to weight and integrate information across the top-*K* candidate peptides for each query protein.

During generation, we optionally expand the retrieved candidate pool to top-*M*peptides (*M* ≥ *K*) and randomly subsample *K* peptides multiple times to construct several stochastic local subgraphs per protein. This yields multiple topology-conditioned vectors that correspond to different retrieved-context views.

### Peptide generator

We generate peptides over an amino-acid vocabulary *A* = {20 canonical residues} augmented with special tokens: *V* = *A* ∪ {BOS, EOS, PAD, UNK, MASK}, where BOS and EOS delimit sequences, PAD denotes padding, UNK denotes out-of-vocabulary residues, and MASK is used only by the auxiliary masked objective. For a peptide *s* = (*s*_1_, …, *s*_*L*_), we construct teacher-forcing inputs and targets as

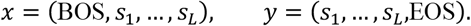

We use a Transformer decoder with *N*_*D*_ layers (default 4) and *H*_*D*_ attention heads (default 8). The model width of the decoder is *d*_model_ (default 512) and it contains dropout (default rate 0.1). The topology-conditioned vector of query protein mentioned above, prot_topo ∈ ℝ^*D*^ is projected into the decoder width and injected as a single memory token:

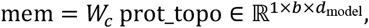

where *b* is the mini-batch size. Token embeddings and learned positional embeddings are summed as *u*_*t*_ = *E*(*x*_*t*_) + *P*(*t*) for position *t* followed by dropout. The decoder computes hidden states *o*_*t*_ and outputs vocabulary logits:

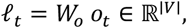

For autoregressive decoding, we apply a standard causal self-attention mask so that each position attends only to ≤ *t*. We use teacher forcing with cross-entropy over all non-padding positions for primary training:

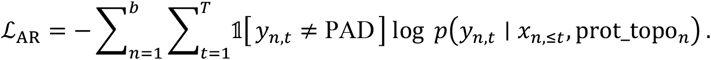

To encourage global consistency while preserving the autoregressive interface, we add an auxiliary Span-MLM objective. For each input *x*, we sample a set of masked positions *M* ⊆ {1, …, *T*} by selecting contiguous spans until reaching an expected masked fraction *p*_mlm_ (default 0.15), excluding BOS and any PAD positions. Span lengths *L*_*s*_ are drawn from a geometric distribution with mean *μ* (default 3) and truncated at *L*_max_ (default 10). At masked positions, we apply a BERT-style replacement rule: 80% replaced by MASK, 10% replaced by a random amino acid, and 10% left unchanged, yielding a corrupted input 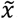 We define MLM labels 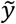 as

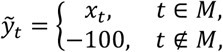

where −100 corresponds to ignored targets in the cross-entropy implementation.

A naïve MLM pass would allow bidirectional attention for all tokens, which encourages reliance on “future” context and thus degrade left-to-right generation. We therefore use a selective bidirectional mask: unmasked query positions remain causal, while masked query positions may attend to the full context. Formally, for a query position *t*, allowed keys *u* would satisfy:

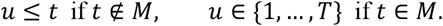

Equivalently, the boolean attention mask *A* ∈ {0,1}^*T*×*T*^ (1 indicates forbidden) is:

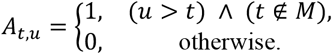

The Span-MLM forward pass uses the same conditioning memory while replacing the standard causal mask with *A*, and we compute cross-entropy only at masked positions as following:

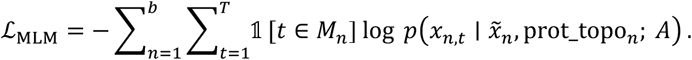

The final objective combines autoregressive and auxiliary losses:

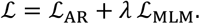

To avoid early optimization instability, we linearly warm up the MLM weight during the first *E*_*w*_ epochs:

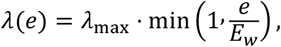

where *e* is the epoch index, and the default of *λ*_max_ is 0.2 and 10 for *E*_*w*_.

In practical conditional generation, a single protein condition may admit multiple valid peptide binders. We therefore perform multi-candidate stochastic decoding per query protein. Given a conditioned vector “prot_topo”, we generate *N* (default 8) peptide candidates by sampling from the autoregressive factorization. At each decoding step, we control diversity by sampling from a temperature-scaled next-token distribution after truncating low-probability residues, while restricting candidates to valid amino-acid tokens (including UNK when enabled) and EOS. To reduce duplicated candidates, we apply a lightweight presence penalty that down-weights tokens already appearing in the partial sequence. We finally deduplicate candidates and keep the top-*N* unique sequences by model score. Each generated sequence *s* is assigned a confidence score computed as its length-normalized log-likelihood under the same model:

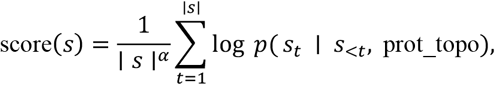

where *α* is a length-normalization exponent (default *α* = 1). The score provides a consistent ranking across candidates and is used for selection and downstream analysis.

### Training procedure

Training is performed on paired protein–peptide examples using teacher forcing. In each mini-batch of size *b*, we first transfer precomputed ESM protein token embeddings **X**^(prot)^ and masks **M**^(prot)^ for the selected protein indices from CPU to GPU. We then encode them with the protein encoder to obtain batch protein representations prot_init ∈ ℝ^*b*×*D*^. Next, using the frozen dual-encoder retriever, we compute query retrieval embeddings for the batch proteins and retrieve a context set of top-K candidate peptides per protein by maximum-inner-product search against the CPU-resident peptide library, yielding corresponding indices top_idx.

We then gather the retrieved peptide token embeddings and masks for top_idx from CPU to GPU and encode them to obtain candidate peptide representations pep_init_sub ∈ ℝ^*b*×*K*×*D*^. These protein and peptide embeddings are fused to a topology-conditioned protein embedding prot_topo ∈ ℝ^*b*×*D*^, which would serve as the conditioning vector for sequence generation.

During peptide generation training, the Transformer decoder receives prot_topo as a single memory token and produces corresponding peptide vocabulary logits. As described before, we use a primary autoregressive cross-entropy objective plus an auxiliary span-masked language modelling (Span-MLM) objective with selective bidirectional attention for masked queries. We use AdamW with learning rate 2 × 10^−4^, mini-batch size 16, and gradient clipping to a global norm of 1.0 for optimization.

We hold out a validation set and use it to monitor training progress. Since peptide generation is inherently multi-solution under a fixed protein condition, we design two complementary evaluation views: (i) teacher-forcing metrics that reflect likelihood modelling and optimization stability, and (ii) free-generation metrics that reflect multi-candidate conditional generation.

For teacher-forcing, we compute token-level negative log-likelihood (NLL) over all non-PAD positions:

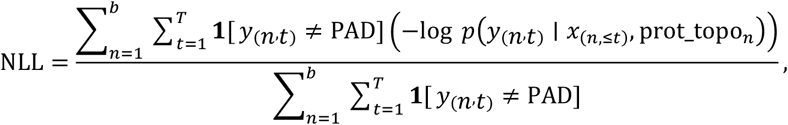

where *b* denotes the mini-batch size, *T* is the padded target length (including EOS), *n* is sample index and *t* is token position. We additionally report perplexity PPL = exp (NLL) and token-level top-5 accuracy over non-PAD positions:

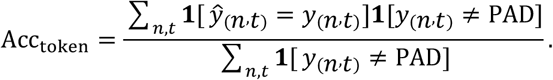

To diagnose over-confident distributions that can lead to mode collapse in generation, we also report the average predictive entropy across non-PAD positions:

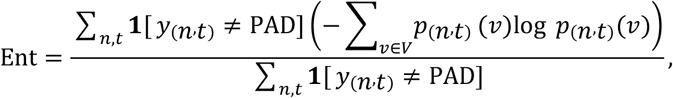

where *p*_(_*n,t*_)_(*v*) is the probability assigned to token *v* at position *t* for sample *n*. We also report the average target length to contextualize these token-level statistics.

For free-generation, we aggregate all associated ground-truth peptides into a multi-reference set 𝒮^∗^(*n*) for each protein in the validation set. We then generate *N* candidate peptides per protein by stochastic autoregressive decoding conditioned on prot_topo_*n*_. Each generated candidate *s* is assigned a confidence score given by its length-normalized log-likelihood:

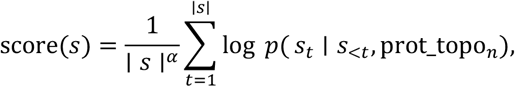

where ∣ *s* ∣ is the generated length for peptide sequence *s, s*_<*t*_ denotes the prefix before position *t*, and *α* is a length-normalization exponent used to make scores comparable across different lengths. Candidates are then ranked by this score. We report multi-reference metrics including Hit@*N* (whether any of the *N* candidates exactly match a reference peptide) and MRR@*N* (the reciprocal rank of the first matching candidate under score-based ranking). We additionally report generation diversity and quality statistics such as uniqueness rate (fraction of distinct sequences among the *N* candidates) and novelty (fraction of generated candidates not appearing in the training set).

## Acknowledgements

I thank my newborn daughter, Anne, whose tiny face repeatedly pulled me back from the brink of giving up. I also thank Dr. Hua Lin, Dr. Chenlin Lu and Mr. Hao Chen for helpful discussions.

## Supplementary Figures

**Figure S1.**
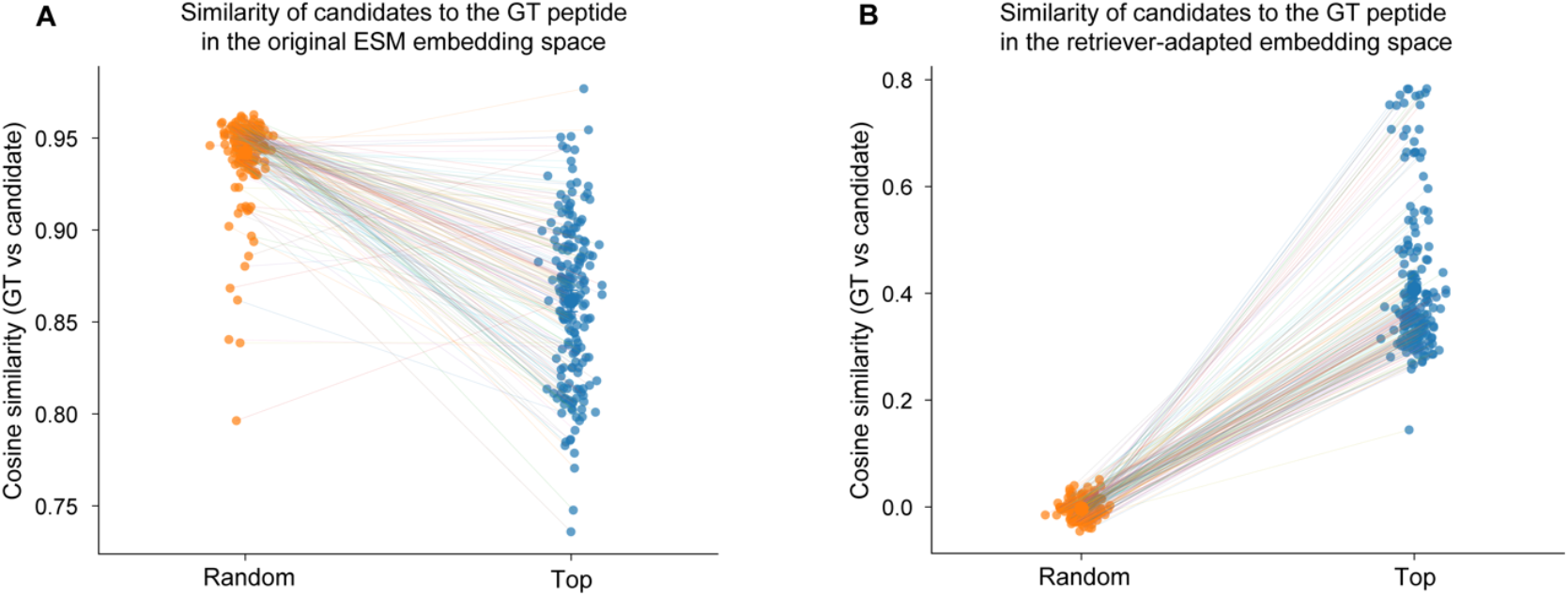
Similarity of candidate peptides to the ground-truth (GT) peptide in the original ESM (A) and retriever-adapted embedding spaces (B). Each dot represents one query protein (n=200). Lines connect paired Random and Top values for the same query.

**Figure S2.**
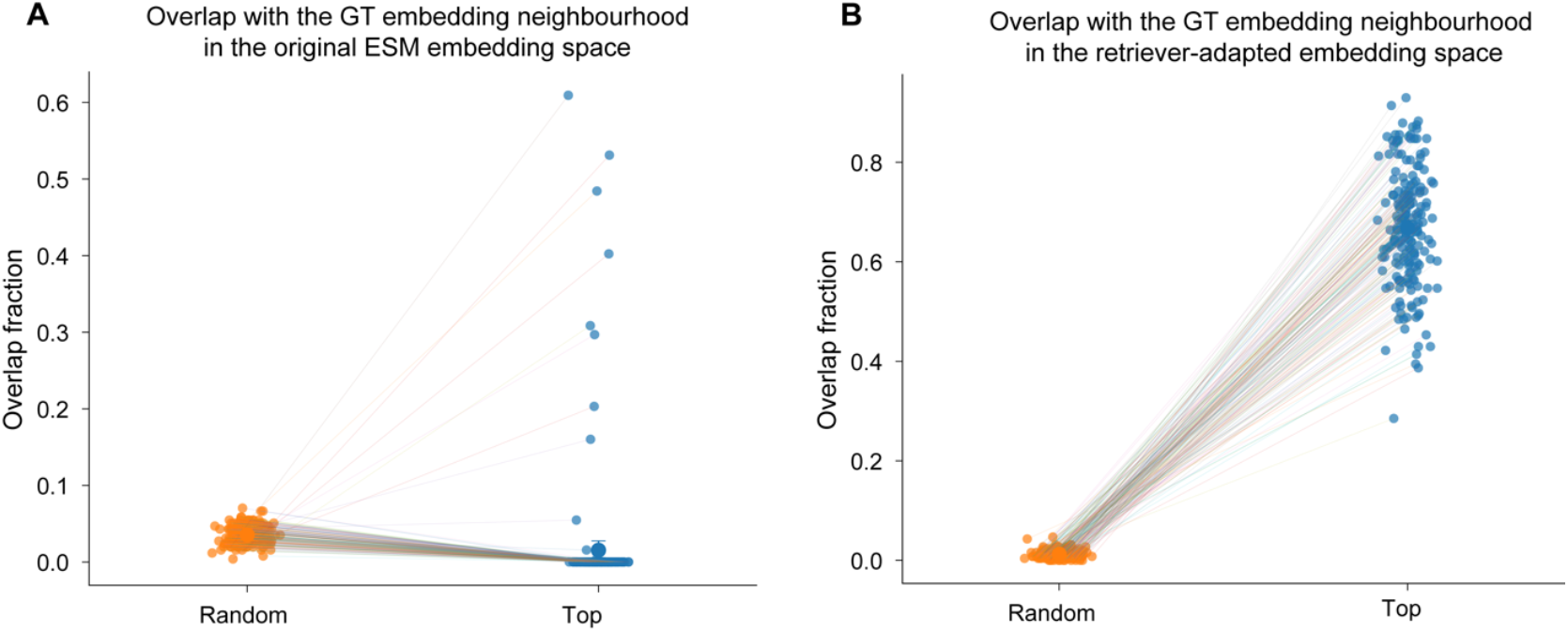
Overlap between retrieved candidates and the ground-truth (GT) peptide neighbourhood in the original ESM (A) and retriever-adapted embedding spaces (B). Each dot represents one query protein (n=200). Lines connect paired Random and Top values for the same query.

**Figure S3.**
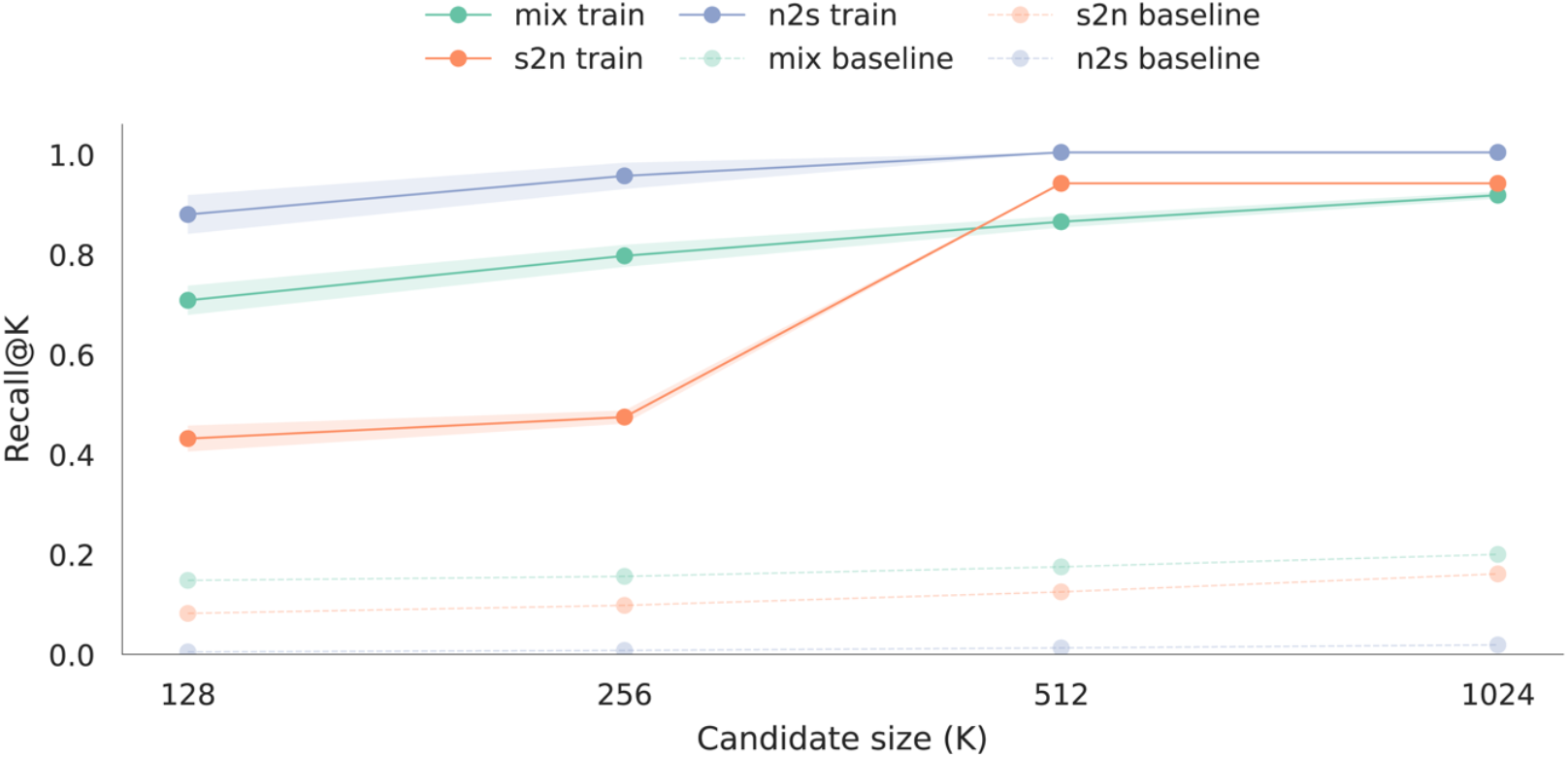
Retrieval performance as a function of candidate-set size. Recall@K is plotted against retrieval depth (K = 128, 256, 512 and 1,024) for our retriever (solid lines) and the ESM baseline (dashed lines) under three evaluation protocols: n2s (train on noisy, test on strict), s2n (train on strict, test on noisy) and mix (noisy + strict with cross-validation). Points indicate the mean across 10 repeats and shaded regions denote s.d.

**Figure S4.**
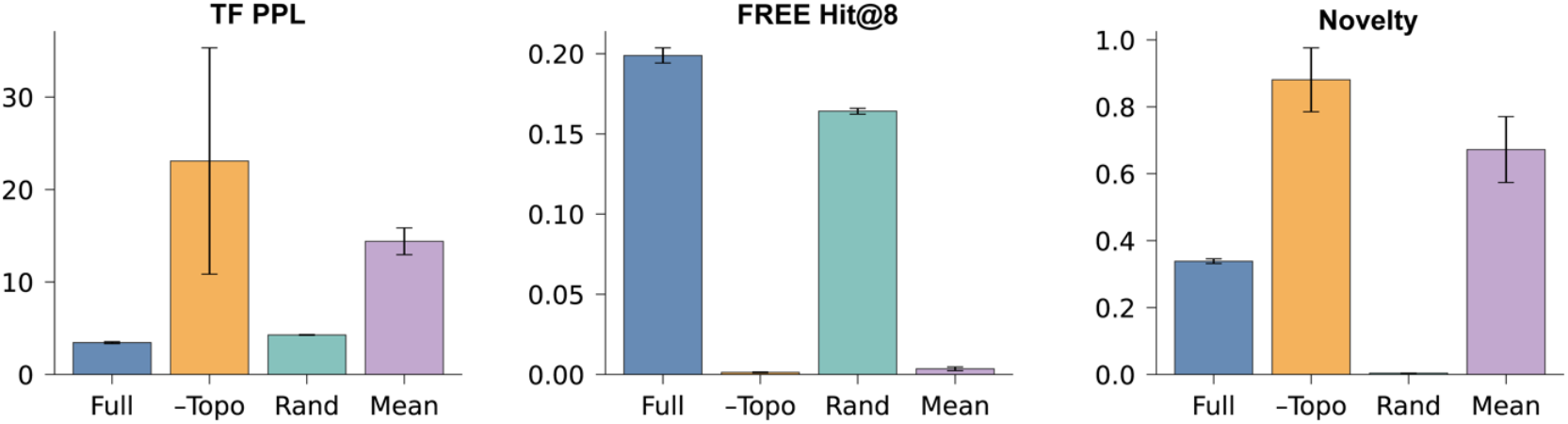
Ablation of topology conditioning using best epochs. Teacher-forcing perplexity (TF PPL), free-generation hit rate among the top 8 generated peptides (FREE Hit@8) and novelty for the full model (“Full”) versus removing the topology conditioner (“–Topo”), using a random selected context instead of the retrieved one (“Rand”) or using mean-pooling in place of the topology conditioner (“Mean”). All comparisons use identical training and decoding settings, and metrics are reported using the best training epoch for each variant. Error bars indicate variability across independent runs.

## Supplementary Tables

**Table S1.** Locality metric summary comparing the original ESM embedding space and the retriever-adapted embedding space. Values are reported as mean ± s.d. across n = 200 independent randomly sampled query targets. For each query, “top” denotes the top-256 retrieved peptides, whereas “rand” denotes 256 peptides sampled uniformly at random from the same whole peptide library. All values are rounded to four decimal places for consistent presentation and interpretation is unchanged under alternative rounding schemes.

| Metric | The original ESM embedding space<br>(mean ± SD) | The retriever-adapted embedding space<br>(mean ± SD) |
| --- | --- | --- |
| IntraSim_top | 0.9102 ± 0.0365 | 0.2589 ± 0.1526 |
| IntraSim_rand | 0.9349 ± 0.0058 | 0.0117 ± 0.0036 |
| GtSim_top | 0.8615 ± 0.0422 | 0.4080 ± 0.1334 |
| GtSim_rand | 0.9421 ± 0.0213 | -0.0034 ± 0.0162 |
| Overlap_top_vs_gtKNN | 0.0154 ± 0.0793 | 0.6679 ± 0.1143 |
| Overlap_rand_vs_gtKNN | 0.0353 ± 0.0114 | 0.0124 ± 0.0078 |
| Seq_intra_JacSim_top | 0.0509 ± 0.0672 | 0.2062 ± 0.2295 |
| Seq_intra_JacSim_rand | 0.0033 ± 0.0025 | 0.0030 ± 0.0019 |
| Seq_gt_JacSim_top | 0.0034 ± 0.0171 | 0.2288 ± 0.2311 |
| Seq_gt_JacSim_rand | 0.0048 ± 0.0101 | 0.0016 ± 0.0027 |
| Seq_intra_BlosumSim_top | -0.0856 ± 0.0870 | 0.0596 ± 0.3251 |
| Seq_intra_BlosumSim_rand | -0.2838 ± 0.0251 | -0.2811 ± 0.0237 |
| Seq_gt_BlosumSim_top | -0.2729 ± 0.0632 | 0.0956 ± 0.3289 |
| Seq_gt_BlosumSim_rand | -0.3003 ± 0.0820 | -0.3066 ± 0.0832 |

**Definitions and computation:** All metrics are computed per query and then averaged across queries. “IntraSim” represents within-set similarity, computed as the mean pairwise similarity among peptides within the “top” set or “rand” set in the corresponding representation space. “GtSim” means similarity to the ground-truth, computed as the mean similarity between peptides in the “top” set or “rand” set and the corresponding ground-truth peptide. “Overlap\_vs\_gtKNN” represents set overlap between the “top” set or “rand” set and the ground truth peptide’s KNN set. “Seq\_intra\_JacSim” and “Seq\_gt\_JacSim” stand for sequence-level Jaccard similarity computed on peptide sequences under the same “intra-set” and “vs ground-truth” definitions as above. “Seq\_intra\_BlosumSim” and “Seq\_gt\_BlosumSim” represent sequence-level similarity computed using a BLOSUM-based scoring scheme under the same “intra-set” and “vs ground-truth” definitions as above. Negative values here indicate lower similarity.

**Table S2.** Paired statistical tests and confidence intervals for Top vs Random in the original ESM embedding space. We report (a) paired effect sizes and two-sided paired significance tests. (b) reports 95% confidence intervals (CIs) for the mean metric values in each condition.

**(a) Paired tests**
| Metric | n | Mean (Top-K) | Mean (Random) | Cohen's d | p (Wilcoxon) | p (t-test) |
| --- | --- | --- | --- | --- | --- | --- |
| IntraSim | 200 | 0.9102 | 0.9349 | -0.625 | 1.97e-16 | 5.06e-16 |
| GtSim | 200 | 0.8615 | 0.9421 | -1.729 | 3.43e-33 | 7.50e-62 |
| Overlap | 200 | 0.0154 | 0.0353 | -0.243 | 3.98e-25 | 7.16e-04 |
| Seq_intra_JacSim | 200 | 0.0509 | 0.0033 | 0.707 | 1.44e-34 | 2.58e-19 |
| Seq_gt_JacSim | 200 | 0.0034 | 0.0048 | -0.070 | 1.29e-02 | 3.25e-01 |
| Seq_intra_BlosumSim | 200 | -0.0856 | -0.2838 | 2.164 | 1.44e-34 | 3.37e-77 |
| Seq_gt_BlosumSim | 200 | -0.2729 | -0.3003 | 0.491 | 9.78e-12 | 5.20e-11 |

**(b) Confidence intervals**
| Metric | 95% CI (Top-K) | 95% CI (Random) |
| --- | --- | --- |
| IntraSim | [0.9050, 0.9149] | [0.9341, 0.9357] |
| GtSim | [0.8557, 0.8673] | [0.9389, 0.9448] |
| Overlap | [0.0057, 0.0275] | [0.0337, 0.0370] |
| Seq_intra_JacSim | [0.0430, 0.0614] | [0.0030, 0.0036] |
| Seq_gt_JacSim | [0.0015, 0.0060] | [0.0035, 0.0063] |
| Seq_intra_BlosumSim | [-0.0960, -0.0717] | [-0.2873, -0.2803] |
| Seq_gt_BlosumSim | [-0.2814, -0.2643] | [-0.3125, -0.2896] |

**Table S3.** Paired statistical tests and confidence intervals for Top vs Random in the retriever-adapted embedding space. We report (a) paired effect sizes and two-sided paired significance tests. (b) reports 95% confidence intervals (CIs) for the mean metric values in each condition.

**(a) Paired tests**
| **Metric** | **n** | **Mean**<br>**(Top-K)** | **Mean**<br>**(Random)** | **Cohen's d** | **p (Wilcoxon)** | **p (t-test)** |
| --- | --- | --- | --- | --- | --- | --- |
| IntraSim | 200 | 0.2589 | 0.0117 | 1.616 | 1.44e-34 | 1.50e-57 |
| GtSim | 200 | 0.4080 | -0.0034 | 3.105 | 1.44e-34 | 2.41e-104 |
| Overlap | 200 | 0.6679 | 0.0124 | 5.482 | 1.43e-34 | 1.21e-150 |
| Seq_intra_JacSim | 200 | 0.2062 | 0.0030 | 0.883 | 1.44e-34 | 8.22e-27 |
| Seq_gt_JacSim | 200 | 0.2288 | 0.0016 | 0.982 | 3.56e-34 | 4.09e-31 |
| Seq_intra_BlosumSim | 200 | 0.0596 | -0.2811 | 1.044 | 2.76e-31 | 8.64e-34 |
| Seq_gt_BlosumSim | 200 | 0.0956 | -0.3066 | 1.312 | 1.85e-34 | 2.83e-45 |

**(b) Confidence intervals**
| **Metric** | **95% CI (Top-K)** | **95% CI (Random)** |
| --- | --- | --- |
| IntraSim | [0.2387, 0.2805] | [0.0111, 0.0122] |
| GtSim | [0.3898, 0.4271] | [-0.0057, -0.0012] |
| Overlap | [0.6515, 0.6828] | [0.0113, 0.0135] |
| Seq_intra_JacSim | [0.1765, 0.2381] | [0.0027, 0.0033] |
| Seq_gt_JacSim | [0.1981, 0.2624] | [0.0013, 0.0020] |
| Seq_intra_BlosumSim | [0.0136, 0.1049] | [-0.2845, -0.2778] |
| Seq_gt_BlosumSim | [0.0516, 0.1404] | [-0.3195, -0.2959] |

**Table S4.** Ablation results at the end of training (70 epochs). We report (a) teacher-forcing and (b) free-generation performance for the full generator (“Full”) and three controlled ablations: “Rand” (retrieved top-K context peptides replaced by randomly sampled peptides), “–Topo” (topology conditioner removed), and “Mean” (retrieved peptides aggregated by mean pooling instead of topology bipartite modulation). Five multiple random seeds are used, and we average across seeds. Values are mean ± s.d. All settings use the same decoder architecture, training recipe, retrieval library, and identical decoding hyperparameters with the same context budget (K=256 peptides) per target; only the specified component is altered in each ablation. Metrics are computed on the same evaluation split. Uniqueness and Novelty summarize diversity and novelty of the top-8 generated peptides, and Soft Hit@8 / Soft MRR@8 provide a similarity-tolerant complement to the strict metrics using the same candidate set. Arrows indicate the preferred direction: (↓) stands for lower is better, whereas (↑) means higher is better.

**(a) Teacher-forcing metrics**
| Setting | TF PPL↓ | TF Top-5↑ |
| --- | --- | --- |
| Full | 3.452 ± 0.104 | 0.9052 ± 0.0003 |
| Rand | 4.056 ± 0.071 | 0.8746 ± 0.0005 |
| -Topo | 352.47 ± 42.41 | 0.2992 ± 0.0016 |
| Mean | 542.12 ± 65.90 | 0.2901 ± 0.0020 |

**(b) Free-generation metrics**
| Setting | FREE Hit@8↑ | FREE MRR@8↑ | Uniqueness↑ | Novelty↑ | Soft Hit@8↑ | Soft MRR@8↑ |
| --- | --- | --- | --- | --- | --- | --- |
| Full | 0.1989 ± 0.0047 | 0.1110 ± 0.0067 | 0.8600 ± 0.0226 | 0.3386 ± 0.0070 | 0.3057 ± 0.0065 | 0.1771 ± 0.0071 |
| Rand | 0.1410 ± 0.0023 | 0.0927 ± 0.0021 | 0.9376 ± 0.0044 | 0.0086 ± 0.0018 | 0.2362 ± 0.0028 | 0.1540 ± 0.0025 |
| -Topo | 0.0000 ± 0.0000 | 0.0000 ± 0.0000 | 0.9504 ± 0.0040 | 1.0000 ± 0.0000 | 0.0000 ± 0.0000 | 0.0000 ± 0.0000 |
| Mean | 0.0000 ± 0.0000 | 0.0000 ± 0.0000 | 0.9398 ± 0.0099 | 0.9994 ± 0.0005 | 0.0000 ± 0.0000 | 0.0000 ± 0.0000 |

**Table S5.**
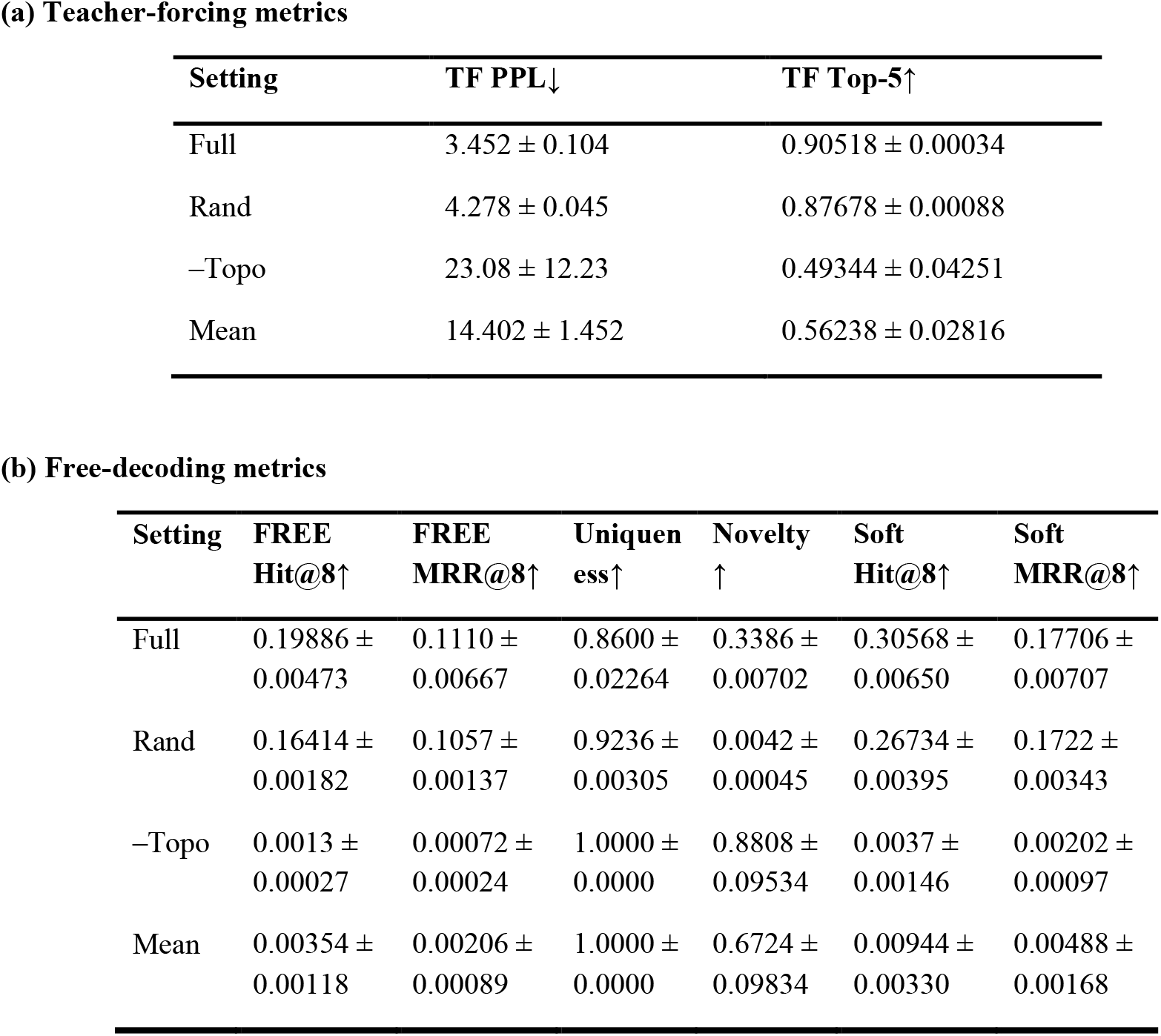
Ablation results at the best checkpoints. Same as Table S1, five multiple random seeds are used, and we average across seeds. Values are mean ± s.d. All settings use the same decoder architecture, training recipe, retrieval library, and identical decoding hyperparameters with the same context budget (K=256 peptides) per target; only the specified component is altered in each ablation.

## References

1. Shen, F. & Dassama, L.M.K. Opportunities and challenges of protein-based targeted protein degradation. Chem Sci 14, 8433–8447 (2023).

2. Bhat, S. et al. De novo design of peptide binders to conformationally diverse targets with contrastive language modeling. Sci Adv 11, eadr8638 (2025).

3. Chen, L.T. et al. Target sequence-conditioned design of peptide binders using masked language modeling. Nat Biotechnol (2025).

4. Chen, T., Hong, L., Yudistyra, V., Vincoff, S. & Chatterjee, P. Generative design of therapeutics that bind and modulate protein states. Current Opinion in Biomedical Engineering 28, 100496 (2023).

5. Dang, C.V., Reddy, E.P., Shokat, K.M. & Soucek, L. Drugging the ‘undruggable’ cancer targets. Nat Rev Cancer 17, 502–508 (2017).

6. Behan, F.M. et al. Prioritization of cancer therapeutic targets using CRISPR-Cas9 screens. Nature 568, 511–516 (2019).

7. Xie, X. et al. Recent advances in targeting the “undruggable” proteins: from drug discovery to clinical trials. Signal Transduct Target Ther 8, 335 (2023).

8. Gao, H., Sun, X. & Rao, Y. PROTAC Technology: Opportunities and Challenges. ACS Med Chem Lett 11, 237–240 (2020).

9. Wang, H. et al. Peptide-based inhibitors of protein-protein interactions: biophysical, structural and cellular consequences of introducing a constraint. Chem Sci 12, 5977–5993 (2021).

10. Bekes, M., Langley, D.R. & Crews, C.M. PROTAC targeted protein degraders: the past is prologue. Nat Rev Drug Discov 21, 181–200 (2022).

11. Liu, X. & Ciulli, A. Proximity-Based Modalities for Biology and Medicine. ACS Cent Sci 9, 1269–1284 (2023).

12. Styles, M.J. et al. PANCS-Binders: a rapid, high-throughput binder discovery platform. Nature Methods 22, 1720–1730 (2025).

13. Diamante, L., Gatti-Lafranconi, P., Schaerli, Y. & Hollfelder, F. In vitro affinity screening of protein and peptide binders by megavalent bead surface display. Protein Eng Des Sel 26, 713–724 (2013).

14. Nielsen, J.C. et al. Machine-Learning-Guided Peptide Drug Discovery: Development of GLP-1 Receptor Agonists with Improved Drug Properties. J Med Chem 67, 11814–11826 (2024).

15. Chen, Z., Wang, R., Guo, J. & Wang, X. The role and future prospects of artificial intelligence algorithms in peptide drug development. Biomed Pharmacother 175, 116709 (2024).

16. Alfaleh, M.A. et al. Phage Display Derived Monoclonal Antibodies: From Bench to Bedside. Front Immunol 11, 1986 (2020).

17. Linciano, S. et al. Screening macrocyclic peptide libraries by yeast display allows control of selection process and affinity ranking. Nat Commun 16, 5367 (2025).

18. Watson, J.L. et al. De novo design of protein structure and function with RFdiffusion. Nature 620, 1089–1100 (2023).

19. Bennett, N.R. et al. Improving de novo protein binder design with deep learning. Nat Commun 14, 2625 (2023).

20. Goto, Y. & Suga, H. The RaPID Platform for the Discovery of Pseudo-Natural Macrocyclic Peptides. Acc Chem Res 54, 3604–3617 (2021).

21. Tsaban, T. et al. Harnessing protein folding neural networks for peptide-protein docking. Nat Commun 13, 176 (2022).

22. Pacesa, M. et al. One-shot design of functional protein binders with BindCraft. Nature 646, 483–492 (2025).

23. Gainza, P. et al. De novo design of protein interactions with learned surface fingerprints. Nature 617, 176–184 (2023).

24. London, N., Raveh, B., Cohen, E., Fathi, G. & Schueler-Furman, O. Rosetta FlexPepDock web server--high resolution modeling of peptide-protein interactions. Nucleic Acids Res 39, W249–253 (2011).

25. Alam, N. et al. High-resolution global peptide-protein docking using fragments-based PIPER-FlexPepDock. PLoS Comput Biol 13, e1005905 (2017).

26. Hosseinzadeh, P. et al. Anchor extension: a structure-guided approach to design cyclic peptides targeting enzyme active sites. Nat Commun 12, 3384 (2021).

27. Chen, S. et al. Design of target specific peptide inhibitors using generative deep learning and molecular dynamics simulations. Nat Commun 15, 1611 (2024).

28. Ding, W. & Gong, H. Predicting the real-valued distances between residue pairs for proteins. arXiv preprint arXiv:1912.06306 (2019).

29. Ding, W. et al. SAMF: a self-adaptive protein modeling framework. Bioinformatics 37, 4075–4082 (2021).

30. Rives, A. et al. Biological structure and function emerge from scaling unsupervised learning to 250 million protein sequences. Proc Natl Acad Sci U S A 118 (2021).

31. Lin, Z. et al. Evolutionary-scale prediction of atomic-level protein structure with a language model. Science 379, 1123–1130 (2023).

32. Hayes, T. et al. Simulating 500 million years of evolution with a language model. Science 387, 850–858 (2025).

33. Ferruz, N., Schmidt, S. & Hocker, B. ProtGPT2 is a deep unsupervised language model for protein design. Nat Commun 13, 4348 (2022).

34. Hie, B.L. et al. Efficient evolution of human antibodies from general protein language models. Nat Biotechnol 42, 275–283 (2024).

35. Madani, A. et al. Large language models generate functional protein sequences across diverse families. Nat Biotechnol 41, 1099–1106 (2023).

36. Brixi, G. et al. SaLT&PepPr is an interface-predicting language model for designing peptide-guided protein degraders. Commun Biol 6, 1081 (2023).

37. Shah, A., Guntuboina, C. & Farimani, A.B. Peptide-GPT: Generative Design of Peptides using Generative Pre-trained Transformers and Bio-informatic Supervision. arXiv preprint arXiv:2410.19222 (2024).

38. Liang, P.-Y., Duran, T. & Bai, J. PepEDiff: Zero-Shot Peptide Binder Design via Protein Embedding Diffusion. arXiv preprint arXiv:2601.13327 (2026).

39. Chen, T. et al. moPPIt: De Novo Generation of Motif-Specific and Functionally Active Peptide Binders via Discrete Flow Matching. bioRxiv, 2024.2007.2031.606098 (2026).

